# Impact of whole-genome duplications on structural variant evolution in the plant genus *Cochlearia*

**DOI:** 10.1101/2023.09.29.560073

**Authors:** Tuomas Hämälä, Christopher Moore, Laura Cowan, Matthew Carlile, David Gopaulchan, Marie K. Brandrud, Siri Birkeland, Matthew Loose, Filip Kolář, Marcus A. Koch, Levi Yant

## Abstract

Polyploidy, the result of whole-genome duplication (WGD), is a major driver of eukaryote evolution. All angiosperms have undergone ancient WGDs, and today stable polyploids account for a substantial portion of both wild and domesticated plant species. Despite its centrality in evolution, WGD is a hugely disruptive mutation, and we still lack a clear understanding of its fitness consequences. Here, we study whether WGD results in greater diversity of genomic structural variants (SVs) and how this influences evolutionary dynamics in nature. Using a set of long-read sequenced samples from the plant genus *Cochlearia* (Brassicaceae), which contains diploids and a recent ploidy series up to octoploid, we show that masking of recessive mutations due to WGD has led to a substantial and progressive accumulation of genic SVs across four ploidy levels. Such SVs likely constitute a genetic load and thus reduce the adaptive potential of polyploid populations. However, this SV accumulation also provides a rich pool of standing genetic variation upon which selection may act in novel environments. By constructing a graph-based pangenome for *Cochlearia*, we identify SVs in hundreds of samples and study the genomic basis of environmental adaptation. We find putatively beneficial SVs involved in pathogen resistance, root development, and salt tolerance, many of which are unique to polyploids. Finally, we explore the adaptive landscapes of SVs and SNPs, identify geographical regions where SVs make novel contributions to adaptive variation, and predict that the role of SVs in environmental adaptation increases due to rapid climate change.

## Introduction

Whole-genome duplication (WGD) is a dramatic mutation that directly challenges the stability of meiosis and DNA management (Bomblies, 2023; Yant and Bomblies, 2015). As such, WGDs are often fatal, but the resulting polyploids that survive these initial obstacles may ultimately thrive (Comai, 2005). Indeed, WGDs have likely contributed to the emergence of all major eukaryotic lineages (Van de Peer et al., 2017), with particular importance in the evolution of plants (Wood et al., 2009). WGDs also have a direct economic impact, as the majority of our most important crop species are polyploid (Salman-Minkov et al., 2016). Understanding how evolutionary dynamics are altered by WGDs is therefore a fundamental goal in evolutionary biology, with applications reaching into agriculture. However, much of the genomic work related to WGDs is conducted on allopolyploids (polyploids resulting from the joining of two lineages), in which the effects of WGDs are confounded by hybridisation. Polyploids resulting from within-species WGDs (autopolyploids), by contrast, provide feasible systems to directly assess how evolutionary processes are shaped by WGDs.

Autopolyploidy is typically characterised by random pairing of chromosomes in meiosis (in allopolyploids chromosome pairing typically happens within subgenomes), resulting in predictable changes in population genetic processes (Haldane, 1930; Wright, 1938). All else being equal, doubling of the genome increases the mutational input, number of recombination events per individual, and the effective population size, leading to an increase in genetic diversity and a decrease in effective linkage (Caballero, 1994; Otto and Gerstein, 2008). Dominance structure is also transformed by WGDs, leading to more efficient masking of recessive mutations (Haldane, 1932; Otto and Whitton, 2000). Thus, increased diversity combined with masking of deleterious mutations may initially raise the adaptive potential of nascent polyploids (Otto, 2007). In the long-term, however, the hidden deleterious mutations might prove difficult to purge and allelic masking not only increases genetic load but also reduces the efficacy of positive selection (Haldane, 1932; Hill, 1970; Otto and Whitton, 2000; Ronfort, 1999), resulting in negative fitness consequences for aging polyploids (Baduel et al., 2018). We can therefore expect both beneficial and detrimental effects arising from WGDs, with empirical support found for some of the theoretical predictions (Baduel et al., 2019; Fisher et al., 2018; Monnahan et al., 2019; Selmecki et al., 2015).

Despite decades of work on polyploid genetics, an aspect that has remained largely unexplored is genomic structural variants (SVs). SVs encompass variants that influence the presence, abundance, location, and/or the orientation of the nucleotide sequence, typically defined as being greater than 50 bp in length. Studies of diploid organisms have established that SVs generally cover much more of the genome than point mutations (Hufford et al., 2021; Liao et al., 2023; Liu et al., 2020; Qin et al., 2021), suggesting that they can have a major impact on the adaptive potential of populations and species. Given their disruptive effects on chromosomal structure, newly emerged SVs tend to be strongly deleterious and thus reduce the fitness of the host (Hämälä et al., 2021; Zhou et al., 2019). Yet SVs have also been associated with adaptive phenotypes in multiple species (Hof et al., 2016; Küpper et al., 2015; Studer et al., 2011; Sutton et al., 2007; Todesco et al., 2020), demonstrating that individual SVs can have beneficial fitness effects. In polyploids, however, the trajectory of SV evolution remains poorly understood, with existing knowledge primarily coming from allopolyploid crop genomes (He et al., 2021; Lovell et al., 2021; Walkowiak et al., 2020). In turn, we are missing an assessment of SV diversity in wild autopolyploid systems, leaving unknown the impact of WGDs on SV evolution in natural contexts.

Here, we study the impact of WGDs on SV landscapes as a function of ploidy, with a focus on understanding the influence of SVs on the adaptive potential of wild autopolyploid populations and species. Given the increased mutational input, combined with more complicated recombination and DNA repair machinery (Bomblies, 2023), we may expect SV emergence to increase in polyploids compared to diploids. We therefore ask how SVs influence the genetic load of polyploid populations, but also explore whether SVs provide unique benefits to polyploids. By analysing hundreds of genomes from the plant genus *Cochlearia*, we find both negative and positive interactions between WGDs and SVs. Masking of recessive mutations has increased the accumulation of deleterious SVs in polyploids, likely reducing the adaptive potential of these populations. However, we also discover apparent benefits resulting from the accumulation of SVs, as many more ploidy-specific SVs contribute to local adaptation in polyploids than in diploids. Finally, we propose that range-edge populations can especially benefit from large-effect SVs, and that SV-mediated adaptation could become more prominent in the future due to rapid climate change. Overall, our results provide important insights into the evolutionary relationship between WGDs and SVs – an aspect that likely has a major impact on the adaptive potential of polyploid organisms.

## Results

### Genetic composition of the *Cochlearia* genus

To study the impact of WGDs on SVs evolution in wild species, we conducted extensive long- and short-read sequencing on the plant genus *Cochlearia* (Brassicaceae). As an evolutionarily dynamic genus, *Cochlearia* exhibits highly labile genome structure, with two base chromosome numbers (*n* = 6 and *n* = 7) and multiple ploidy levels (from diploids to dodecaploids) found among the species (Crane and Gairdner, 1923; Gill, 1971; Koch et al., 1998, 1996; Koch, 2012; Saunte, 1955). Here, we focus on populations from the diploid *n* = 6 species *C. pyrenaica*, *C. excelsa, C. aestuaria*, and *C. islandica*; diploid *n* = 7 species *C. groenlandica* and *C. triactylites*; tetraploid *n* = 6 species *C. officinalis, C. micacea*, and *C. alpina*; hexaploid species *C*. *tatrae*, *C. bavarica*, and *C. polonica*; and octoploid species *C. anglica*. The tetraploids likely resulted from within-species WGDs (autopolyploids), as evidenced by widespread multivalent formation at meiosis (Bray et al., 2023), whereas the evolutionary history of the higher ploidies is more complex, potentially involving both auto- and allopolyploidisation events. See Koch (2012) and Wolf et al. (2021) for comprehensive discussions of the evolutionary history of the *Cochlearia* species.

In total, our dataset comprised 23 samples sequenced with Oxford Nanopore (ONT) or Pacific Biosciences (PacBio) long-read technologies and 351 samples sequenced with Illumina short-read technology. The individuals represent 76 populations, covering the primary range of *Cochlearia* throughout Europe (Fig. 1A), along with locations in Svalbard and North America (Dataset S1). We first used SNP data derived from short-read sequencing to examine patterns of genetic diversity and differentiation among the *Cochlearia* populations. Compared to the diploid populations, polyploids exhibited lower levels of genetic diversity (Fig. 1B) and more negative Tajima’s *D* (Fig. 1C), potentially reflecting bottlenecks and subsequent expansions resulting from the recent establishment of these populations (Wolf et al., 2021). A principal component analysis (PCA) indicated genetic clustering both due to ploidy and geographical location: the first principal component (PC) primarily corresponded to a difference between diploids and polyploids, whereas the second PC revealed clustering among multiple locations (Fig. 1D). The geographical clustering was also evident in within-ploidy PCAs, while little separation was found based on species assignments (Fig. S1). We further discovered a signal of isolation-by-distance, with between population *F*_ST_ estimates increasing with geographical distance, especially among the diploids (Fig. 1E). However, by using redundancy analysis (RDA) to model the role of geography, climate, and ploidy in explaining differentiation among the populations, we found climatic conditions to be a better predictor of genetic differentiation than either geographical distance or ploidy (Fig. 1F).

**Figure 1.**
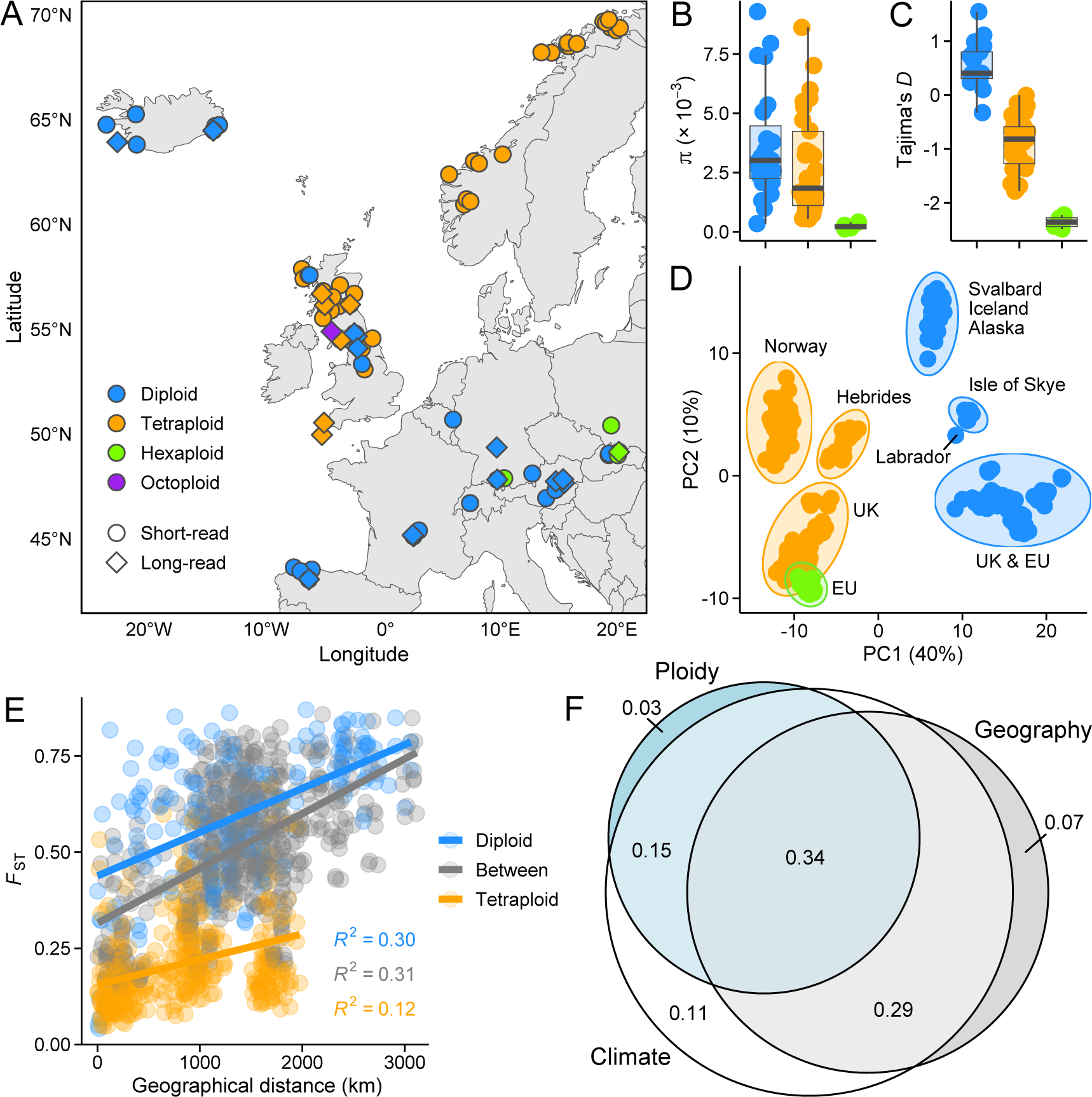
Locations and genetic variation among *Cochlearia* populations in this study. **A**: Map depicting European sampling locations. Shown are short-read sequenced populations (circles) and long-read sequenced individuals (diamonds). **B**: Pairwise nucleotide diversity (π) estimates for the short-read sequenced populations. **C**: Tajima’s *D* estimates for the short-read sequenced populations. **D**: First two axes of a PCA. The proportion of variance explained by the PCs is shown in parentheses. **E**: Relationship between *F*_ST_ and geographical distance among diploid and tetraploid populations (between = diploid vs. tetraploid). **F**: The role of geography, climate, and ploidy in explaining genetic differentiation among these *Cochlearia* populations. Adjusted *R*^2^ values from partial RDA models are shown in the circles. Note that the same colour legend applies to panels A–E.

### SV identification and methylation assessment using long-read sequencing

Based on the analysis of SNP data, we found evidence that polyploidy influences the genetic composition of the *Cochlearia* genus. To explore whether WGDs also have an impact on SV landscapes, we performed long-read sequencing to identify SVs in 23 samples chosen to represent diverse lineages and ploidies. However, due to low sequencing depth, we excluded four diploids from our main analyses (Table S1), resulting in a set of 10 diploids, seven tetraploids, one hexaploid, and one octoploid. After aligning reads against the chromosome build *C. excelsa* reference genome (Bray et al., 2023), we used Sniffles2 (Smolka et al., 2022) to identify SVs from the alignments. First, as Sniffles2 was developed primarily for diploid organisms, we used simulated data to confirm that it likely has a good power to detect SVs in our high-depth (mean depth = 68) autotetraploid samples (Fig. 2A and Table S2). We focused our analyses on insertions and deletions between 50 and 100,000 bp in size and filtered them for variant quality, missing data, and sequencing depth. After filtering, we retained 33,376 SVs in diploids and 47,755 in tetraploids. As both sequencing depth and read length can influence the power to detect SVs, we confirmed that tetraploids also carried more SVs after downsampling the alignments to equal number of base pairs (Fig. S2; *P* = 0.001, Wilcoxon rank-sum test). By comparing the SV sequence against our transposable element (TE) library, we found that in both diploids and tetraploids ∼45% of the SVs contained TE sequence (Fig. S3), suggesting that many of the SVs are likely the result of TE mobilisation. To examine whether TE activity, and thus the potential of TEs to generate new SVs, differs between the ploidies, we quantified TE methylation using our ONT sequenced samples. Although we observed higher methylation levels in tetraploids than in diploids (Fig. S4), the pattern was not unique to TEs, and once we controlled for the genome-wide difference in methylation levels, we saw no evidence that TE families are systematically hyper- or hypomethylated in tetraploids (Fig. S5). Indeed, by estimating putative insertion times for TEs, we found no significant differences between the ploidies (Fig. S6), indicating that differential silencing of TEs is not a major factor shaping SV landscapes in diploids and tetraploids.

**Figure 2.**
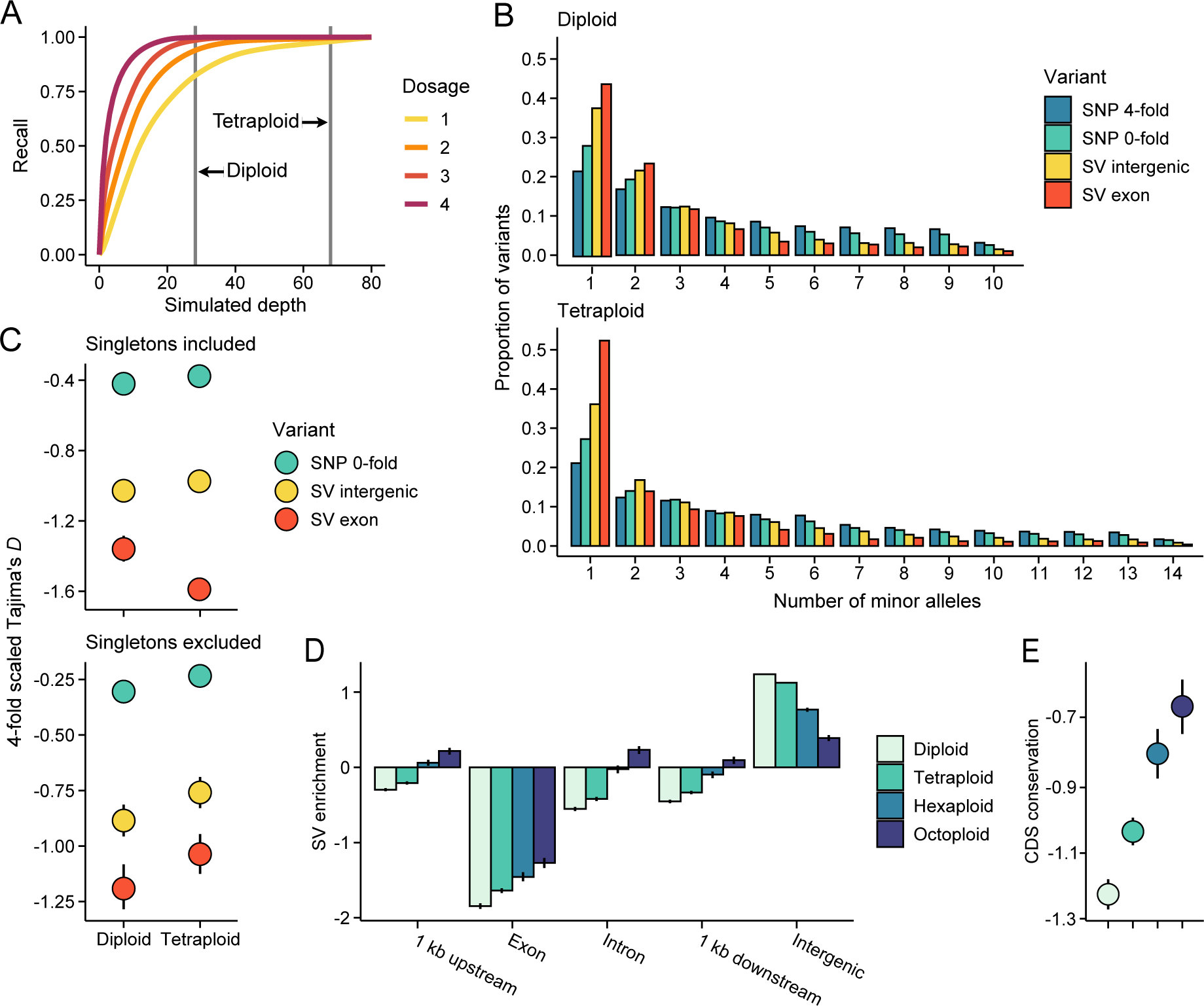
Fitness effects of structural variants (SVs). **A**: Validation of the SV caller. Shown are recall estimates for different levels of read depth over simulated insertions and deletions. For each of the simulated scenarios, precision estimates were > 0.95. Grey horizontal lines mark the mean sequencing depth of our diploid and tetraploid samples. **B**: Folded allele frequency spectra (AFS) for different variant types in diploids and tetraploids. **C**: Tajima’s *D* estimated from the whole AFS (singletons included) and AFS with singletons excluded. Values for nonsynonymous (0-fold) SNPs and SVs were scaled in relation to synonymous (4-fold) sites. **D**: SV enrichment across different genomic features in diploids and polyploids. Shown are log_2_-transformed ratios of observed to expected numbers of SVs. **E**: Coding sequence (CDS) conservation of genes affected by SVs (exon in panel D). Shown are median standardised GERP scores estimated among 30 eudicot species (lower values indicate weaker conservation). Note that the same colour legend applies to panels D and E. In panels C, D, and E, error bars indicate 95% bootstrap-based CIs.

### Masking progressively increases SV load in polyploids

To gain insight into the fitness effects of SVs, we estimated allele frequency spectra (AFS) for SVs found at genic (exons) and intergenic (> 5 kb away from genes) regions and compared these to nonsynonymous (0-fold) and synonymous (4-fold) SNPs. Overall, we found the AFS of diploids and tetraploids to be similar, except for an excess of rare exonic SVs in tetraploids (Fig. 2B). In the absence of mutation rate difference, such excess could either indicate stronger purifying selection in tetraploids or that recently emerged SVs are tolerated at functional regions because their effects are being masked by the additional allelic copies. To answer this question, we summarised the AFS using Tajima’s *D* and compared two sets of estimates: ones estimated from the whole AFS and ones with singletons (i.e., variants with only single allele present) excluded. We found that the exclusion of singletons removed the excess of rare exonic SVs (Fig. 2C), supporting the idea that such SVs are being retained due to more efficient masking in tetraploids (as stronger purifying selection would skew the whole AFS towards rare variants). Therefore, our AFS-based analyses suggest that masking of recessive mutations allows SVs to accumulate in tetraploids that would have been purged by purifying selection in diploids. We acknowledge, however, that these analyses rely on the correct identification of the SV genotypes, which can be challenging in polyploids, despite our validation (Fig. 2A). Thus, as an alternative approach, we examined the genomic locations of the SVs (regardless of their genotypes) and compared the observed numbers of SVs found overlapping different genomic features to random expectations. Given that this analysis does not require population-level data, we also included the hexaploid and octoploid samples. As expected, we discovered an overall deficit of exonic SVs and an excess of intergenic SVs (Fig. 2D). However, the deficit was greater in diploids than in polyploids, with the amount decreasing progressively with increasing ploidy (Fig. 2D). Furthermore, by examining the level of coding sequence conservation at genes affected by the SVs, we found a similar cline between all ploidies (Fig. 2E), indicating that SVs are being retained in genes under stronger selective constraint in polyploids than in diploids. Both results further support our conclusion that masking allows recessive SVs to accumulate in polyploids, likely progressively increasing the genetic load of higher ploidy populations.

### *Cochlearia* pangenome reveals the influence of SVs in environmental adaptation

Although our analyses of the long-read data suggest that masking has increased the accumulation of deleterious SVs in polyploids, we might expect that some SVs provide selective benefits for the *Cochlearia* populations. Therefore, to examine the role of SVs in environmental adaptation, we constructed a graph-based pangenome for *Cochlearia* and used it to genotype 135,574 SVs in 351 short-read sequenced samples. After filtering the SVs for variant quality, missing data, and minor allele frequency (MAF), we used 11,393 (diploids), 18,313 (tetraploids), and 15,959 (all: diploids, tetraploids, and hexaploids) SVs to conduct genotype-environment association (GEA) analyses. Our analysis identified 49 SVs strongly associated with climatic variables in diploids, 159 in tetraploids, and 122 when considering all ploidy levels (Fig. 3A). Among the top outliers, we discovered SVs closely adjacent (< 1 kb) to genes *ACT7* (diploids), involved in root development and germination (Gilliland et al., 2003); *RIN2* (tetraploids), involved in pathogen resistance (Kawasaki et al., 2005); and *SIGA* (all), involved in control of plastid gene expression (Onda et al., 2008). Gene ontology (GO) terms related to root development and pathogen resistance (hypersensitive response) were also highly enriched among all outliers in diploids and tetraploids, respectively (Fig. 3B). By combining the three outlier sets, we discovered enrichment for additional GO terms, such as cellular response to salt (Fig. 3B). The candidate genes and biological processes associated with outlier SVs were largely independent of those identified with SNPs, as fewer than expected genes (odds ratio = 0.02; *P* < 2 × 10^-16^, Fisher’s exact test) and none of the GO terms were shared between the SV- and SNP-based outlier sets.

**Figure 3.**
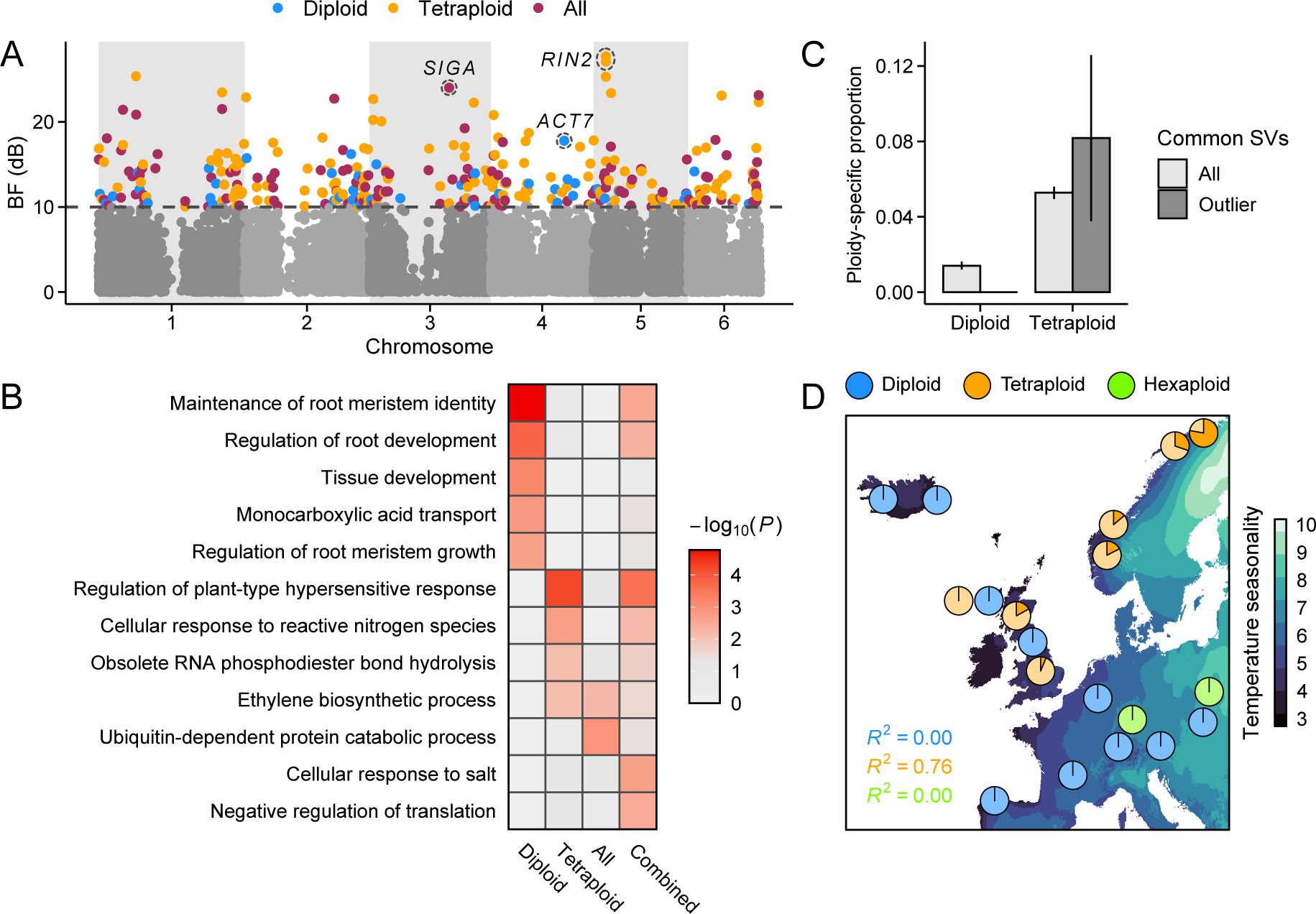
Footprints of environmental adaptation at SVs. **A**: Bayes Factor (BF) estimates from genotype-environment association (GEA) analyses conducted among diploid, tetraploid, and all (diploids, tetraploids, and hexaploids) populations. SVs with BF (dB) ≥ 10 were considered putatively adaptive. Top candidate gene (< 1 kb of the top outlier SV) from each analysis is highlighted. **B**: Results from a gene ontology (GO) enrichment analysis conducted on candidate genes. Included are GO terms with *P* < 0.01 from each of the three GEA sets as well as from a set with all outliers combined. **C**: The proportion of common SVs (MAF > 0.05) found only among diploid or tetraploid populations (ploidy-specific). Error bars show 95% bootstrap-based CIs. **D**: Example of a tetraploid-specific outlier SV, found 107 bp upstream of a gene *RIN2.* Pie charts show the frequencies of reference (lighter colour) and alternative (darker colour) alleles in closely adjacent populations.

Given the greater accumulation of SVs in polyploids, we might assume that ploidy-specific SVs are more likely to contribute to adaptation in tetraploids than in diploids. Consistent with this expectation, we discovered that larger proportion of common SVs (MAF > 0.05) were ploidy-specific in tetraploids than in diploids (Fig. 3C; *P* < 2 × 10^-16^, Fisher’s exact test), including GEA outlier SVs (Fig. 3C; *P* = 0.01, Fisher’s exact test). For example, two top outlier SVs in our GEA, closely adjacent to gene *RIN2*, were only polymorphic among the tetraploid populations (Fig. 3D). We further note that the proportion of ploidy-specific SVs in tetraploids is likely underestimated, as our long-read data covers only part of the tetraploid range (the large cluster of diversity in Norway is missing, Fig. 1D), whereas among diploids there was a good correspondence between long- and short-read sequencing (Fig. 1A).

### The role of SVs in environmental adaptation increases due to climate change

Our results suggest that SVs make novel contributions to environmental adaptation in *Cochlearia*. To gain more insight into the geographical distribution of this adaptive variation, we predicted adaptive landscapes across the European range of the genus (Fig. S7). By leveraging the associations between genetic and environmental variables, adaptive landscapes can be used to project adaptive variation to unsampled locations (Fitzpatrick and Keller, 2015) and to model population vulnerability under climate change (Rellstab et al., 2021). Here, however, we extend this approach to identify geographical regions where SVs potentially make unique contributions to adaptive variation by visualising the difference between SV- and SNP-based landscapes. Our analysis identified the northern (Norway and Iceland) and southern (Spain and France) edges of the *Cochlearia* range as locations where the adaptive landscapes are most strongly diverged (Fig. 4). Furthermore, by conducting a prediction using environmental variables projected for years 2061 – 2080, we discovered that the disparity between SV- and SNP-based adaptive landscapes may increase due to climate change (assuming that populations mainly track climate change through existing variation), especially in populations at the southern edge of the *Cochlearia* range (Fig. 4). We note that the same populations are not, according to our analysis, the ones most vulnerable to climate change, which are primarily found at the eastern edge of the *Cochlearia* range (Fig. S8).

**Figure 4.**
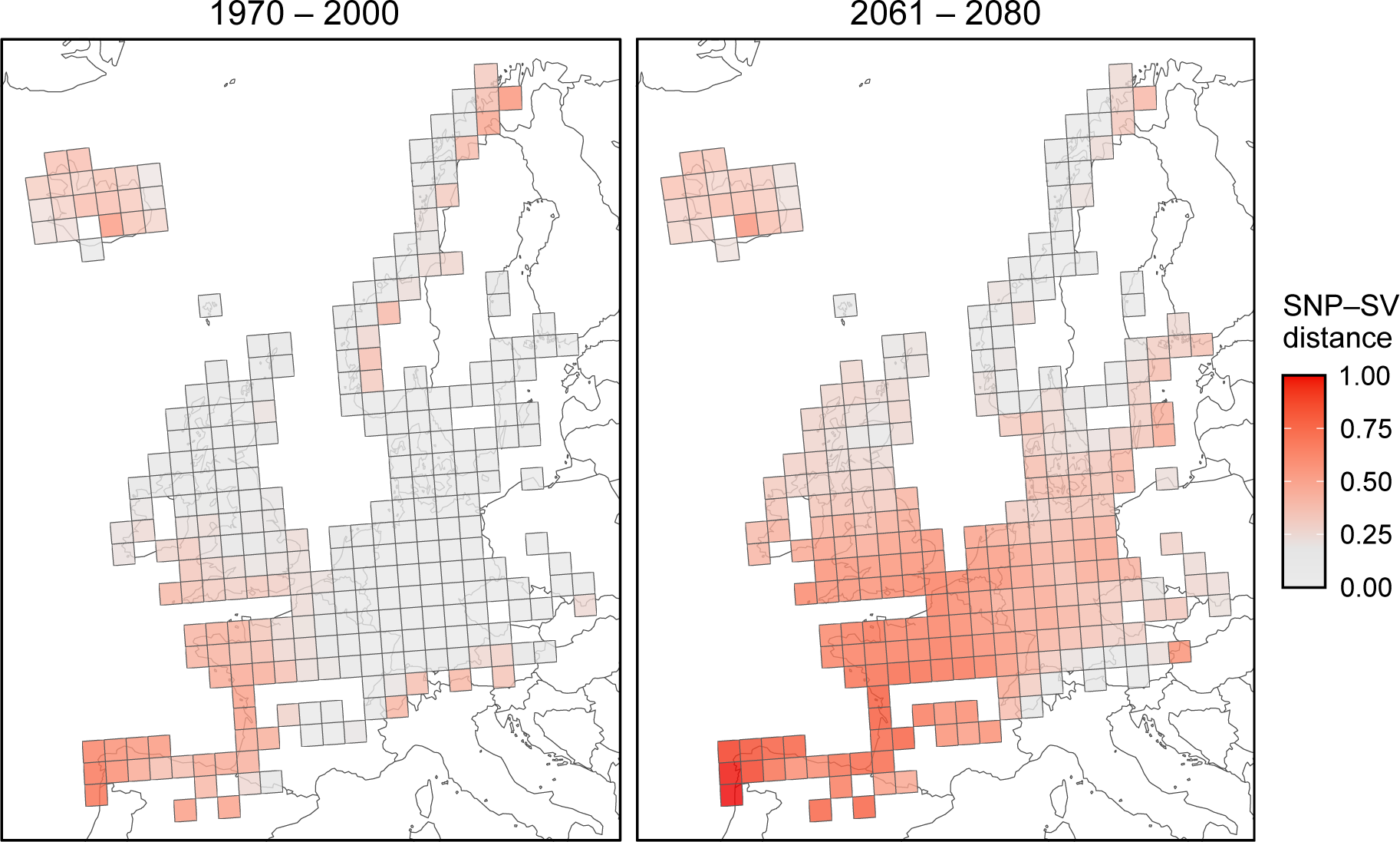
Adaptive distance between SNPs and SVs across the European range of *Cochlearia*. Colour scale indicates the level of unique contributions that SVs are predicted to make to adaptive variation (Euclidean distance between the adaptive indices). First panel shows past adaptation, based on 11 bioclimatic variables collected between 1970 and 2000, and the second panel shows a projection for years 2061 – 2080.

## Discussion

How WGDs influence adaptive evolution is a long-standing question in evolutionary biology. Based on both theoretical (Haldane, 1930; Hill, 1970; Otto and Whitton, 2000; Wright, 1938) and empirical (Baduel et al., 2019; Fisher et al., 2018; Monnahan et al., 2019; Selmecki et al., 2015) work, we can expect pervasive fitness consequences arising from WGDs. However, the evolutionary relationship between WGDs and SVs is poorly understood, partly because SV identification has been challenging using short-read sequencing technologies (Mahmoud et al., 2019).

Here, we used long-read sequencing and pangenomics to study the impact of WGDs on SV landscapes in the plant genus *Cochlearia*. We discovered a substantial accumulation of genic SVs in polyploids that likely would have been purged by purifying selection in diploids. Theory suggests that such hidden load can have a major impact on the long-term fate of polyploid populations (Haldane, 1932; Otto and Whitton, 2000; Ronfort, 1999), contributing to eventual extinction or rediploidisation (Baduel et al., 2018). Although previous studies have discovered an excess of nonsynonymous SNPs (Monnahan et al., 2019) and TEs (Baduel et al., 2019) in recently founded autotetraploids, we may expect that SV accumulation has a particularly strong effect on the genetic load of polyploid populations. SVs are not only more deleterious than point mutations (Hämälä et al., 2021; Zhou et al., 2019) but also could be more frequently generated in polyploids due more complicated recombination and DNA repair machinery (Bomblies, 2023), as experimentally shown in yeast (Selmecki et al., 2015). Indeed, we previously showed that genes involved in DNA repair have evolved rapidly in the tetraploid *C. officinalis* since its divergence from the diploid *C. pyrenaica* (Bray et al., 2023), suggesting that WGD in *Cochlearia* has resulted in a shift in the (internal) selective environment due to extra challenges in DNA management.

Assuming that most deleterious mutations are partially recessive (Agrawal and Whitlock, 2011), SVs could have two major consequences for the fate of autopolyploid populations: 1) The point at which the genetic load of a newly formed autotetraploid population exceeds that of its diploid progenitor is reached faster with stronger selection coefficients (Otto and Whitton, 2000), meaning that SVs (compared to point mutations) could shorten the period of beneficial fitness effects arising from WGDs. 2) Once population reaches equilibrium, the fitness reduction due to deleterious mutations is roughly equal to the product of ploidy level and mutation rate (Otto and Whitton, 2000; Ronfort, 1999), indicating that higher rate of SV emergence in polyploids would increase the genetic load beyond that predicted from ploidy alone. Therefore, the accumulating SV load is likely an important factor limiting the adaptive potential of polyploid organisms, especially among the higher ploidies. We acknowledge, however, that in our hexaploid and octoploid samples the load inference could be influenced by their mixed auto- and allopolyploid histories (Wolf et al., 2021), as subgenome dominance and lack of homoeologous recombination may increase the accumulation of deleterious mutations in allopolyploids compared to autopolyploids (Conover and Wendel, 2022). Nevertheless, the progressive accumulation of genic SVs across the four ploidy levels supports the idea that increasing ploidy leads to more efficient masking of recessive mutations, thus reducing the efficacy of purifying selection.

Despite the increased SV loads in polyploids, we also discovered apparent benefits resulting from the SV accumulation. Ploidy-specific SVs were more likely to contribute to local adaptation in tetraploids than in diploids, suggesting that the greater SV diversity in polyploids occasionally gets harnessed by positive selection. Furthermore, as interploidy gene flow is almost exclusively unidirectional from diploids to tetraploids (Baduel et al., 2018; Morgan et al., 2021), tetraploids are more likely to benefit from adaptive SVs originating in diploids than *vice versa*. Therefore, our results suggest that SVs contribute to the greater diversity of adaptive alleles available for polyploids (Bohutínská et al., 2023), compensating for some of the detrimental effects arising from the increased SV load. By analysing genes closely adjacent to the outlier SVs, we discovered enrichment of genes involved in different biological processes. The most prominent were related to root development in diploids, pathogen resistance in tetraploids, and salt tolerance in the combined dataset. The fast evolving resistance genes have been associated with SVs in multiple plant species (Bayer et al., 2020), highlighting the importance of structural variation in the ongoing arms race between plants and their pathogens (Anderson et al., 2010). Additionally, SVs have been linked to salinity adaptation in *Arabidopsis* (Busoms et al., 2018), making SV-mediated adaptation a potential contributor to the high salt tolerance exhibited by these *Cochlearia* species (Bray et al., 2023; Pegtel, 1999). The *Cochlearia* species (particularly the diploid *C. pyrenaica*) are also highly metallicolous (Nawaz et al., 2017), which could be associated with root development (Dijk et al., 2022) and thus SV-mediated adaptation. Importantly, these candidate genes and biological processes were not detected using SNPs, demonstrating that SVs need to be considered for a comprehensive view of adaptive processes.

To gain more insight into the unique roles of adaptive SVs, we searched for differences between SV- and SNP-based adaptive landscapes (Capblancq and Forester, 2021). The northern and southern range edges were highlighted as regions where the adaptive landscapes are most strongly diverged, potentially indicating novel contributions made by SVs. Indeed, we might expect to find more adaptive SVs in range-edge populations, as large-effect mutations tend to be favoured in populations that are far from their selective optima (Hämälä et al., 2020; Orr, 1998). By conducting a prediction using future climate projections, our modelling further suggests that SVs currently conferring adaptation to the southern environment could increase in frequency and spread northward due to climate change, resulting in greater divergence between the SV- and SNP-based adaptive landscapes. Furthermore, this analysis expects that populations track shifting fitness optima through existing variation, but SV emergence could also increase due to climate change, as environmental stress is known to induce TE mobilisation (Hämälä et al., 2022; Wos et al., 2021), potentially providing more opportunities for SV-mediated adaptation.

### Conclusions

By conducting extensive long- and short-read sequencing on the plant genus *Cochlearia*, we have gained novel insights into the evolutionary relationship between WGDs and SVs. We discovered a progressive accumulation of genic SVs across four ploidy levels, indicating increased SV loads in polyploids compared to diploids. Given the strongly negative fitness effects of SVs, we expect such SV loads to limit the long-term adaptability of polyploid populations and species. However, by constructing a graph-based pangenome for *Cochlearia*, we also found putative benefits arising from the SV accumulation, as ploidy-specific SVs were more likely to contribute to local adaptation in tetraploids than in diploids. Finally, our modelling work highlighted the unique roles of SVs in adaptation to current and future climates. Overall, our comprehensive analysis of SVs in *Cochlearia* sheds light on important and largely unexplored aspect of polyploid genomes, broadening the perspective of polyploid evolution as well as the evolution of structural variation in wild populations and species.

## Material and methods

### High molecular weight DNA isolation, Oxford Nanopore, and PacBio HiFi sequencing

To study the evolutionary role of SVs in *Cochlearia*, we collected samples from 23 individuals to be used in long-read sequencing. The set included 14 diploids, seven tetraploids, one hexaploid, and one octoploid (Table S1). Before starting DNA isolation, Carlson lysis buffer (100 mM Tris-HCl, 2% CTAB, 1.4 M NaCl, 20 mM EDTA, 1% PEG 8000) was mixed with 0.3 g PVPP and 50 ul B-mercaptoethanol and preheated to 65 °C. Leaf material from individual plants was ground into the heated solution and incubated for an hour at 65 °C. 20 ml chloroform was then added and mixed by inverting. The mixture was centrifuged at 3500 rpm (4 °C) for 15 minutes, the top layer of the lysate added to 1 × volume isopropanol, inverted to mix, and incubated at –80 °C for 15 minutes before being centrifuged at 3500 rpm (4°C) for 45 minutes. The supernatant was removed, the pellet air dried (sterile wipes were also used to dry the side walls of the tube) and resuspended in 500 ul nuclease-free water containing 2 ul of RNase A before being left to incubate at 37°C for 45 minutes. Samples were column purified with a Qiagen Blood and Cell Culture DNA Maxi Kit clean up. The DNA concentration was checked on a Qubit Fluorometer 2.0 (Invitrogen) using the Qubit dsDNA HS Assay kit. Fragment sizes were assessed using the Genomic DNA Tapestation assay (Agilent). Removal of short DNA fragments and final purification to high molecular weight DNA was performed with the Circulomics Short Read Eliminator XS kit. After DNA isolation, two samples were used for Pacific Biosciences (PacBio) HiFi sequencing and 21 samples for Oxford Nanopore Technologies (ONT) sequencing.

ONT libraries were prepared using the Genomic DNA Ligation kit SQK-LSK109 following manufacturer’s procedure. Libraries were loaded onto R9.4.1 PromethION Flow Cells and run on a PromethION Beta sequencer. Due to the rapid accumulation of blocked flow cell pores or due to apparent read length anomalies on some *Cochlearia* runs, flow cells used in the runs were treated with a nuclease flush to digest blocking DNA fragments before loading with fresh libraries according to the ONT Nuclease Flush protocol (version NFL_9076_v109_revD_08Oct2018). FAST5 sequences produced by PromethION sequencer were basecalled using the Guppy6 (https://community.nanoporetech.com) high accuracy basecalling model (dna_r9.4.1_450bps_hac.cfg) and the resulting FASTQ files quality filtered by the basecaller. PacBio sequencing was performed on a Sequel IIe at Novogene Europe (Cambridge, UK) in CCS mode.

### Short-read library preparation and sequencing

We also used a set of 109 short-read sequenced *Cochlearia* individuals from Bray et al. (2023), which includes 39 diploids and 70 tetraploids. Although this sampling covers several locations across the Western and Northern Europe, it is mainly focused on the UK. To expand our sampling to more varied environments, we additionally collected 242 *Cochlearia* individuals across Europe and North America, leading to a final set of 351 individuals from 76 populations used for short-read sequencing (Dataset S1).

DNA was prepared using the commercially available DNeasy Plant Mini Kit from Qiagen (Qiagen: 69204). Illumina libraries were constructed from genomic DNA using Illumina DNA Prep library kit and IDT for Illumina DNA/RNA Unique Dual Index sets. Library preparation was performed using a Mosquito HV (SPT Labtech) liquid handling robot. The standard protocol timings and reagents were used, but with 1/10^th^ reagent volumes at all steps. A total of 9 – 48 ng of DNA was used as library input and 5 cycles of PCR were used for the library amplification step. Individual libraries were pooled together and size selected using 0.65 × AMPure XP beads to minimise library fragments < 300 bp. Library pools were sequenced on a Novaseq 6000 using 2 × 150 bp paired-end reads at Novogene Europe (Cambridge, UK).

### Transposable element annotation

We previously identified TEs from the *C. excelsa* reference genome (Bray et al., 2023). However, as the reference originated from a selfing diploid, we additionally assembled the genomes of one outcrossing diploid (*C. pyrenaica*) and one outcrossing tetraploid (*C*. *officinalis*) to expand our library of *Cochlearia* TEs. To do so, the individuals were sequenced using PacBio HiFi reads to an estimated depth of ∼20 (diploid) and ∼40 (tetraploid) × the haploid genome size. The reads were then *de novo* assembled using hifiasm (Cheng et al., 2021) and haplotigs purged from the primarily assemblies using purge_dups (Guan et al., 2020). The resulting assemblies had a total size of 359 (diploid) and 315 (tetraploid) mb, with contig N50 of 2.6 mb (diploid) and 630 kb (tetraploid). BUSCO (Simão et al., 2015) analysis indicated high completeness of the gene space, with 96% of the single-copy Brassicales genes found in both assemblies (Fig. S9). As with the *C. excelsa* reference genome, we annotated the assemblies using the EDTA pipeline (Ou et al., 2019), which includes multiple methods to comprehensively identify both retrotransposons and DNA transposons. To generate a single TE library across the three species, we used the cleanup_nested.pl script from EDTA to remove redundant (> 95% identical) consensus sequences from the combined library. We last conducted BLAST queries against a curated plant protein database from Swiss-Prot to remove likely gene sequences from the TE library. See Fig. S10 for outline of the annotated TE superfamilies.

### Short-read processing and variant calling

Low quality reads and sequencing adapters were removed using Trimmomatic (Bolger et al., 2014) and the surviving reads aligned to the *C*. *excelsa* refence genome using bwa-mem2 (Vasimuddin et al., 2019). Although we aligned reads from multiple species against a single reference, alignment proportions were high for all samples (between 80% and 99%), likely reflecting the shallow divergence between the *Cochlearia* species (Wolf et al., 2021). We removed duplicated reads using Picard tools (https://broadinstitute.github.io/picard/) and identified SNPs using GATK4 (Van der Auwera and O’Connor, 2020). Filtering of the variant calls was based on the GATK’s best practices protocol, and we included filters for mapping quality (MQ ≥ 40 and MQRankSum ≥ –12.5), variant confidence (QD ≥ 2), strand bias (FS < 60), read position bias (ReadPosRankSum ≥ –8), and genotype quality (GQ ≥ 15). We further removed SNPs with per-sample sequencing depth ≥ 1.6 × the mean depth to avoid issues caused by paralogous mapping.

### Analyses of genetic variation

We used the short-read-based SNP calls to infer genetic relationships among our diploid and polyploid *Cochlearia* populations. First, we estimated pairwise nucleotide diversity (π) and Tajima’s *D* for each population using both mono- and biallelic sites. We then conducted a principal components analysis (PCA) using linkage-pruned (*r*^2^ ≤ 0.1 within 100 SNPs, minor allele frequency [MAF] > 0.05) SNPs found at synonymous (4-fold) sites. Following Patterson et al., (2006), we estimated a covariance matrix representing the genetic relationships among each pair of individuals. For two individuals, *i* and *j*, covariance (*C*) was calculated as:

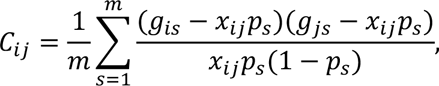

where *m* is the number of variable sites, *g_is_* is the genotype of individual *i* in site *s, x* is the average ploidy level of the two individuals, and *p* is the alternate allele frequency. We then conducted PCA on the matrix using the R function prcomp and extracted the first two axes of the rotated data for plotting. We also estimated genetic differentiation between populations using *F*_ST_. Here, we employed the *F*_ST_ measure by Hudson et al. (1992), as recommended by Bhatia et al. (2013).

To disentangle drivers of genetic differentiation among the *Cochlearia* populations, we tested for a pattern of isolation-by-distance and isolation-by-environment. Following Capblancq and Forester (2021), we performed redundancy analyses (RDA) using the R package vegan (Oksanen et al., 2022). We first estimated allele frequencies for the populations using linkage-pruned SNPs with MAF > 0.05 and ≤ 20% missing data. Missing population frequencies were imputed by randomly drawing them from a beta distribution with scale parameters calculated from the mean and variance of the non-missing values. We then extracted all 19 bioclimatic variables from WorldClim (Fick and Hijmans, 2017) and conducted forward model selection using RDA to identify variables explaining a significant proportion of genetic variation. Based on 1000 permutations, we kept 14 variables with *P* < 0.01. To transform the spatial structure of our data into a format usable in RDA, we conducted a principal coordinates analysis on a geographical distance matrix, retaining ten principal coordinates after forward model selection. Last, using partial RDA, we decomposed the effects of climate, geography, and ploidy in explaining genetic variation among the *Cochlearia* populations.

### Validation of the SV caller

We aligned both ONT and PacBio long-reads against the *C. excelsa* reference genome using minimap2 (Li, 2018) and identified SVs from the alignments using Sniffles2 (Smolka et al., 2022). Our main analyses were based on 10 diploids (we excluded four diploids due to low sequencing depth, Table S1) and seven tetraploids. As Sniffles2 expects the reads to originate from diploid organisms, we first used simulated data to evaluate its performance in autotetraploids. To estimate parameter values for the simulations, we used NanoPlot (De Coster et al., 2018) to calculate the mean and SD of read lengths across all samples. By aligning reads from the *C. excelsa* reference individual against the reference genome, we estimated an empirical error rate of 4% for the ONT reads. We note, however, this is likely a conservative estimate, as we assumed that all differences between the assembly and sequencing reads were due to sequencing errors, whereas such differences may also result from erroneous alignments and heterozygous SNPs (although heterozygous SNPs should be relatively rare in the selfing reference individual). We then randomly chose a single 10 kb region from the reference genome and introduced a 1 kb insertion or deletion into it. Using PBSIM2 (Ono et al., 2021), we generated simulated ONT reads from the modified and unmodified FASTA files, and combined them assuming average read proportions for simplex (1/4), duplex (2/4), triplex (3/4), and quadruplex (4/4) mutations. The total read depth was either 5, 10, 20, 40, or 80 × the simulated region. As Sniffles2 includes a preset for detecting rare SVs (i.e., SVs that are supported by only few reads), we assessed the performance of both the standard and rare SV modes. Based on the results (Table S2), we chose the rare SV mode for all subsequent Sniffles2 runs.

### Long-read-based variant calling and genotyping

We called SV candidates individually for each sample and joined them into multi-sample VCF files using the population calling algorithm in Sniffles2 (Smolka et al., 2022). To reduce false positives caused by erroneous alignments, we included the reference (highly homozygous) individual in all multi-sample VCF files and excluded SVs that were called as heterozygotes or alternate homozygotes in the refence sample. We further focused our analyses on insertions and deletions (variant quality ≥ 20) between 50 bp and 100 kb, as methods based on read alignments are generally less accurate at detecting other types of SVs (e.g., tandem duplications and inversions) as well as very large SVs (Mahmoud et al., 2019). We also identified SNPs from the long-read alignments using Clair3 (Zheng et al., 2022). Running Clair3 on a model trained with R9.4 and Guppy6 reads (or alternatively with HiFi for the two PacBio sequenced individuals), we generated gVCF files for each individual and combined them into multi-sample VCF files using GATK4 (Van der Auwera and O’Connor, 2020).

Although our simulations suggest that Sniffles2 has good power to detect insertions and deletions in autotetraploids (Table S2), the genotype calls are incorrect due to diploid-specific genotyping model. Therefore, we collected allele count data (i.e., the number of reads supporting the reference and alternate alleles in each variant) for SVs and SNPs, and used the R package Updog (Gerard et al., 2018) to estimate genotype likelihoods and probabilities. We required that variants used in Updog had ≤ 20% missing data and were covered by ≥ 10 reads in diploids and ≥ 20 reads in tetraploids. To include genotype uncertainty directly into our analyses, we estimated the allelic dosage, or the expected genotype, from the genotype probabilities as 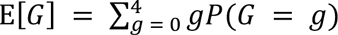, where *G* is the genotype. We then repeated this dosage estimation for the diploids to make the ploidy comparison equal.

### Differential methylation analysis

To quantify DNA methylation, we used Deepsignal-plant (Ni et al., 2021) to detect 5-Methylcytosine (5mC) states of individual bases in the ONT data. To do so, we first used Tombo (Stoiber et al., 2017) to assign basecalls and genomic locations to raw signal reads. Then, using a model trained on *Arabidopsis thaliana* and *Oryza sativa* R9.4 reads, we had Deepsignal-plant estimate methylation frequencies in three sequence contexts, CG, CHG, and CHH (where H is A, T, or C), for each ONT sequenced individual with a mean depth ≥ 10. We last used cytosines covered by ≥ 6 reads to calculate methylation levels across genes and TEs.

We searched for differentially methylated TE families between diploids and tetraploids to assess TE activity. To do so, we used logistic regression and likelihood-ratio tests (LRTs) to identify associations between methylation levels and the ploidy. To control for the effects of population structure on methylation patterns, we conducted PCA on genome-wide methylation levels, used scree plots to define the number of main components in the data, and included the extracted PCs as cofactors in the models. *P*-values from the LRTs were transformed to false discovery rate-based *Q*-values (Storey, 2002) to account for multiple testing. We considered TE families with *Q* < 0.05 as differentially methylated between the ploidies.

### Estimation of TE insertion times

We identified non-refence TE insertions from the ONT alignments using TELR, which has shown good performance in highly heterozygous, polyploid samples (Han et al., 2022). TELR combines Sniffles (we modified the pipeline to use Sniffles2 in the rare SV mode) and RepeatMasker (http://www.repeatmasker.org) to first identify TE insertions and then performs local assembly of the inserted sequences using wtdbg2 (Ruan and Li, 2020). After running TELR on each sample, we aligned the inserted sequences against the consensus TEs using MAFFT (Nakamura et al., 2018) and calculated sequence divergence (*K*) using the F81 substitution model (Felsenstein, 1981) implemented in the R package phangorn (Schliep, 2011). Last, we estimated insertion times as 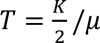, where μ is the per year substitution rate, here assumed to be equal to the per generation mutation rate estimated for *Arabidopsis thaliana* (6.95 × 10^-9^ per base pair, Weng et al., 2019).

### Fitness effects of SVs

We assessed the fitness effects of SVs by first analysing their allele frequency spectra (AFS). Using the estimates of allelic dosage, we calculated folded AFS for SVs found in genic (exons) and intergenic (> 5 kb away from genes) regions and compared them to synonymous (4-fold) and nonsynonymous (0-fold) SNPs. In the case of missing data (max 20%), we imputed the missing alleles by drawing them from a Bernoulli distribution. We further evaluated the selective removal of SVs by calculating the ratio of observed to expected numbers of SVs found overlapping different genomic features (1 kb up- and downstream of genes, exons, introns, intergenic). The expected numbers were estimated by defining the proportion of the genome that is covered by each feature (i.e., under random expectations, SVs would be distributed according to those proportions). We note, however, that these expectations could be affected by variation in mutations rates and insertion preference of TEs, but here were assumed that such biases are, on average, equal between the ploidies (this was confirmed for TEs, Figs. S4 – S6).

To determine the level of selective constraint on genes affected by the SVs, we estimated coding sequence conservation using GERP++ (Davydov et al., 2010). We first selected 29 eudicot species from the clade Superrosidae (Table S3), whose divergence times ranged from 20 million years (*Lobularia maritima*) to 123 million years (*Vitis vinifera*) in relation to *C. excelsa* (Hohmann et al., 2015). To identify sequence homologs, we conducted BLAST searches against species-specific protein databases, selecting only the best match with an *e*-value < 1 × 10^−5^ for each gene. We aligned the coding sequences using MAFFT (Nakamura et al., 2018), keeping only homolog sets with 15 or more species. We then chose 1000 random genes with no missing species, extracted synonymous sites based on the *C. excelsa* sequence, and estimated a maximum likelihood tree using the R package phangorn (Schliep, 2011). Based on the species tree and multiple alignments, we used GERP++ to estimate the rejected substitutions score for sites in the *C. excelsa* coding sequence, indicating the degree of nucleotide conservation relative to the synonymous substitution rate. Finally, we normalised the GERP scores using the range of possible values (as the range depends on the sample size of a particular site), calculated a median for each gene, and standardised the gene-specific estimates to a median of zero and MAD (median absolute deviation) of one.

### Pangenome construction and SV genotyping

To more broadly study the evolutionary impact of SVs in *Cochlearia*, we genotyped our long-read based SVs in a set of 351 short-read sequenced individuals (Dataset S1). First, we identified SVs from all 23 long-read sequenced individuals, including four diploids previously excluded due to low sequencing depth, one hexaploid, and one octoploid, to construct a pangenome graph to serve as a reference for the short-real alignments. We kept all insertions and deletions filling the following requirements: not identified in the reference individual, length between 50 bp and 100 kb, variant quality ≥ 20, supported by ≥ 4 reads, and the proportion of supporting reads ≥ 0.1 of all reads. We then used vg (Garrison et al., 2018) to construct a pan-genome graph based on the chromosome build *C. excelsa* reference genome and the resulting 135,574 SVs. The short-read data were aligned to the pangenome graph using vg map (Garrison et al., 2018) and SVs genotyped using vg call (Hickey et al., 2020). We last combined the individual-based SV calls into multi-sample VCF files using BCFtools (Li, 2011) and estimated genotype probabilities and allelic dosage using Updog (Gerard et al., 2018).

### Genotype-environment association analyses

We tested for an association between genetic and environmental variables to identify loci involved in local adaptation. To do so, we characterised the growing environment of 70 European *Cochlearia* populations (we excluded three populations from North America and three populations from Svalbard, as they represented clear climatic outliers) using 11 bioclimatic variables (Fig. S11) identified with RDA (see “Analyses of genetic variation“ for more details) and conducted genotype environment association (GEA) analyses using BayPass (Gautier, 2015). BayPass was ran on SV and SNP data compiled for three sets of samples: diploids, tetraploids, and all (diploids, tetraploids, and hexaploids combined). Note that BayPass works on population-specific allele frequencies (and not individual genotypes), making it suitable for polyploids. We required the variants to have MAF > 0.05 and ≤ 20% missing data to be included in the analyses. To control for the confounding effects of population structure, we included covariance matrices estimated using synonymous, linkage-pruned SNPs into all BayPass runs. Following the recommendation of Gautier (2015), we repeated each run ten times with different seed numbers (settings for the priors and the MCMC sampling were left default) and calculated a median Bayes Factor (BF) for the variants. Variants with median deciban (dB) BF ≥ 10 were considered putatively adaptive (corresponding to strong evidence for an association between genetic and environmental variables).

To better understand the functional importance of the outlier SVs, we conducted gene ontology (GO) enrichment analyses using the R package topGO (Alexa and Rahnenfuhrer, 2023). For each outlier SV and SNP, we included the closest gene within 1 kb and ran GO enrichment analyses using the weight01 algorithm and Fisher’s exact test. We defined the background distribution of GO terms using only genes ≤ 1 kb of SVs and SNPs. Following the recommendation of Alexa and Rahnenfuhrer (2023), we considered GO terms with *P* < 0.01 as significantly enriched among the candidate gene sets.

### Adaptive landscapes and climate change

We used RDA (Oksanen et al., 2022) to explore the adaptive landscapes of SVs and SNPs. First, we estimated population allele frequencies for each outlier loci identified with BayPass (loci combined from diploid, tetraploid, and all runs) and imputed missing frequencies by drawing them from a beta distribution. We then used RDA to search for multivariate association between allele frequencies and the 11 bioclimatic variables used in the GEA. Following Capblancq and Forester (2021), we used the loadings of the first two RDA axes to predict the adaptive index for each environmental pixel across Europe (see Fig. S12 for a biplot of the loadings). We then acquired occurrence data for the *Cochlearia* genus from Global Biodiversity Information Facility (GBIF.org, 2023), cleaned up the records using both an automated tool (Zizka et al., 2019) and manual curation, and summarised the results into 100 × 100 km grid points comprising the European range of *Cochlearia*. We did this prediction using both outlier SNPs and SVs, and plotted the adaptive distance (Euclidean distance between the adaptive indices) between the two variant types to identify geographical regions where SVs potentially make unique contributions to the adaptive landscape. To explore possible effects of climate change on the adaptive landscapes, we used a Shared Socioeconomic Pathways (SSP) scenario SSP3-7.0 (Yukimoto et al., 2019) to model the increase in greenhouse gas concentrations by years 2061 – 2080.

## Data availability

Sequence data for this study have been deposited in the European Nucleotide Archive (ENA) at EMBL-EBI under accession number PRJEB66308. Scripts for conducting the analyses are available at https://github.com/thamala/polySV.

## Author contributions

L.Y., T.H, and M.A.K. conceived the study. T.H. performed analyses. C.M., L.C., M.C., D.G., and M.L. performed laboratory work. M.A.K., M.K.B., S.B., and F.K. provided materials. T.H. wrote the manuscript with input from other authors.

## Acknowledgements

We thank S. Bray for assistance with sample collection, and J. Brookfield and A. MacColl for comments on the manuscript. We are grateful for access to the University of Nottingham’s Deep Seq sequencing facility and Augusta HPC service. The project has received funding from the European Union’s Horizon 2020 research and innovation programme under the Marie Skłodowska-Curie grant agreement No. 101022295 (to T.H.) and the European Research Council (ERC) grant agreements No. 679056 (to L.Y.) and No. 850852 (to F.K.).

