## Supplementary information for "Impact of whole-genome duplications on structural variant evolution in the plant genus *Cochlearia*"

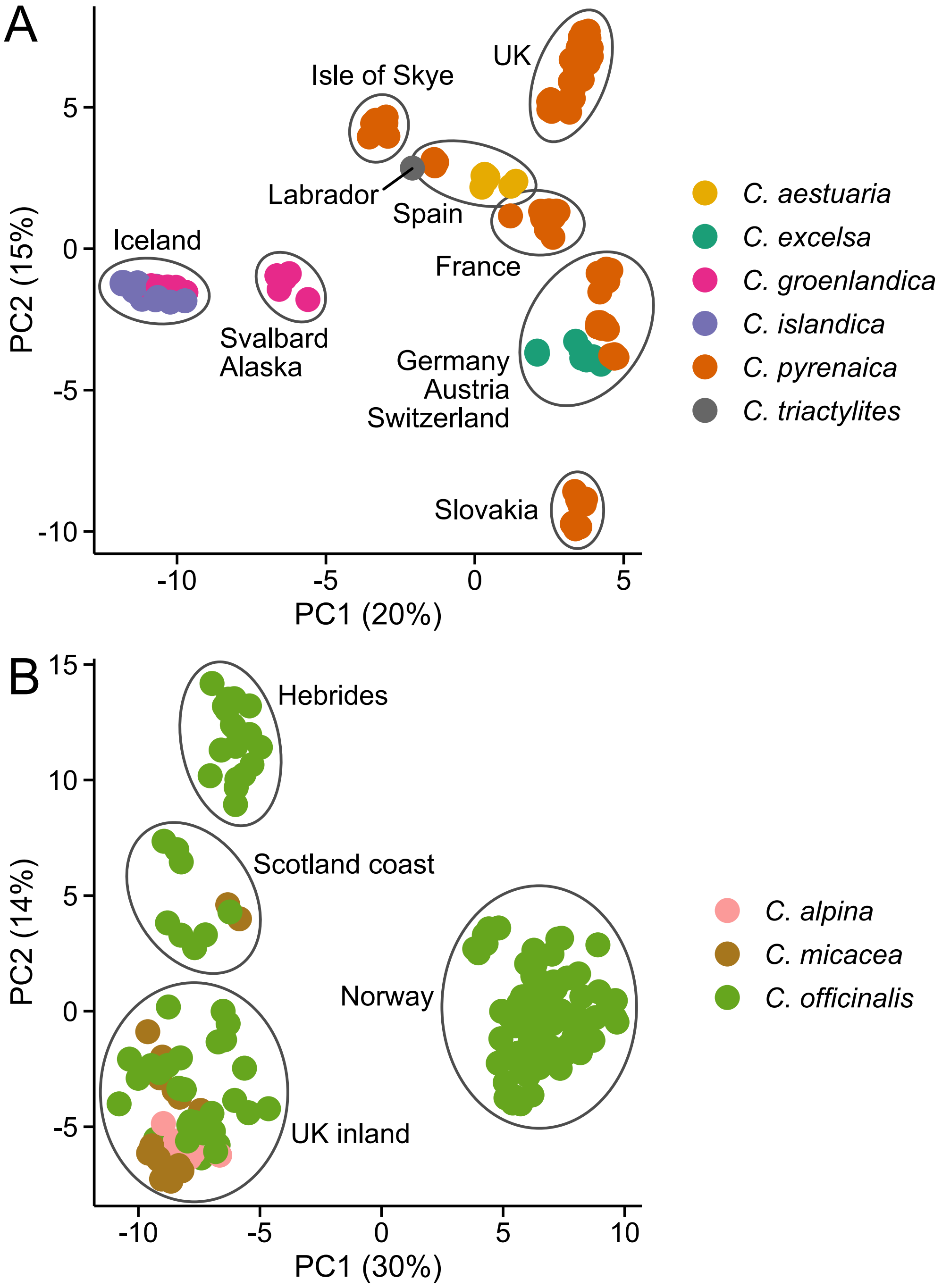


**Figure S1.** PCA conducted separately on diploids (**A**) and tetraploids (**B**). Species are marked with colours. The proportion of variance explained by the PCs is shown in parentheses.

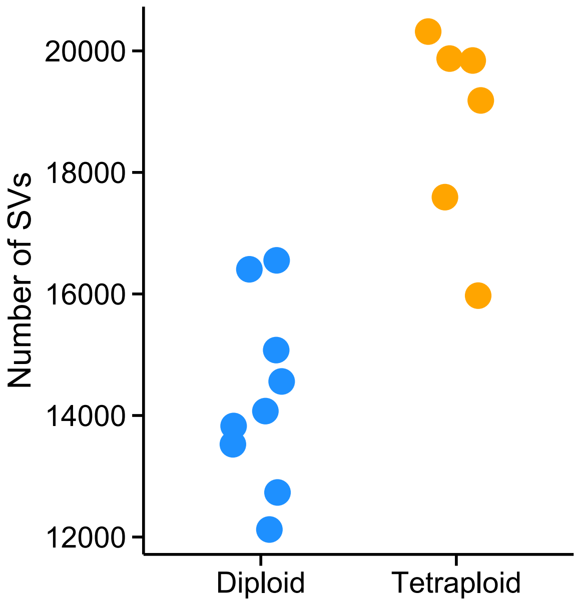


**Figure S2.** Number of SVs found in ONT sequenced samples after equalising the alignments to same number of base pairs covered (mean read length × number of reads).

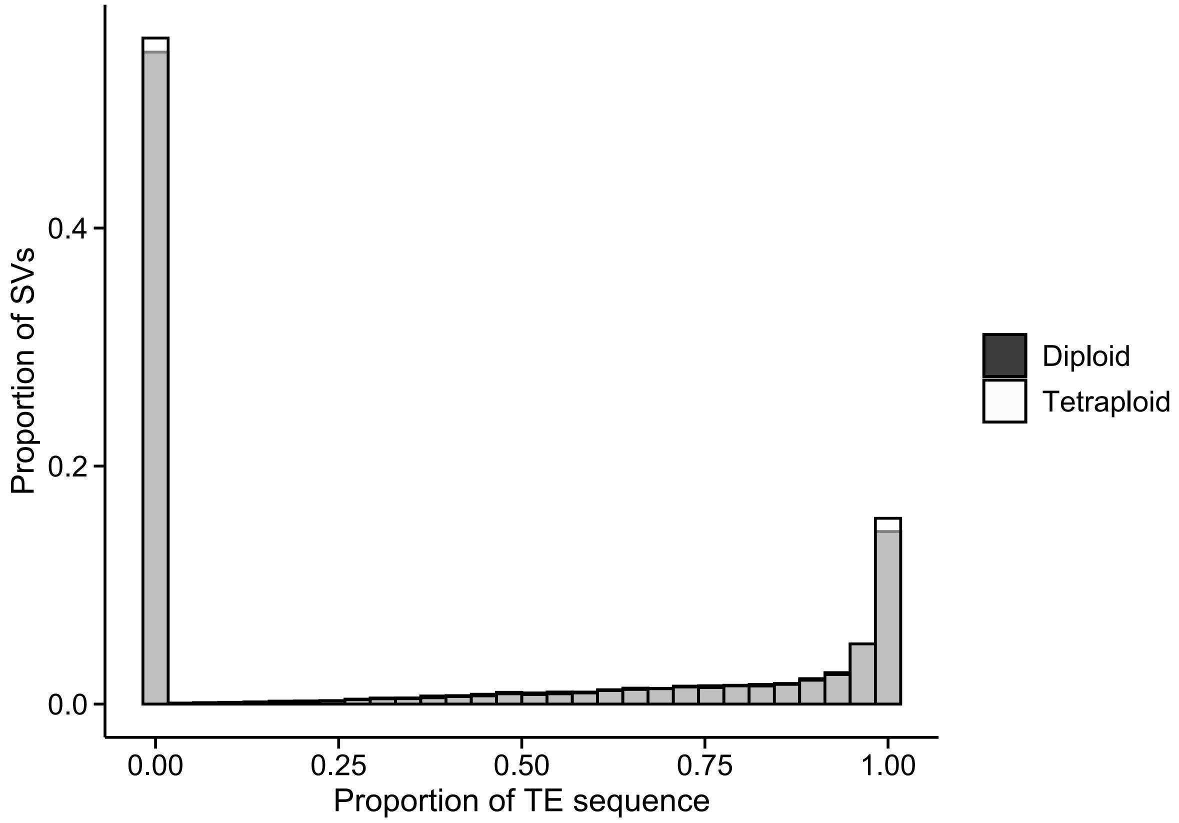


**Figure S3**. Proportion of TE sequence found in SVs.

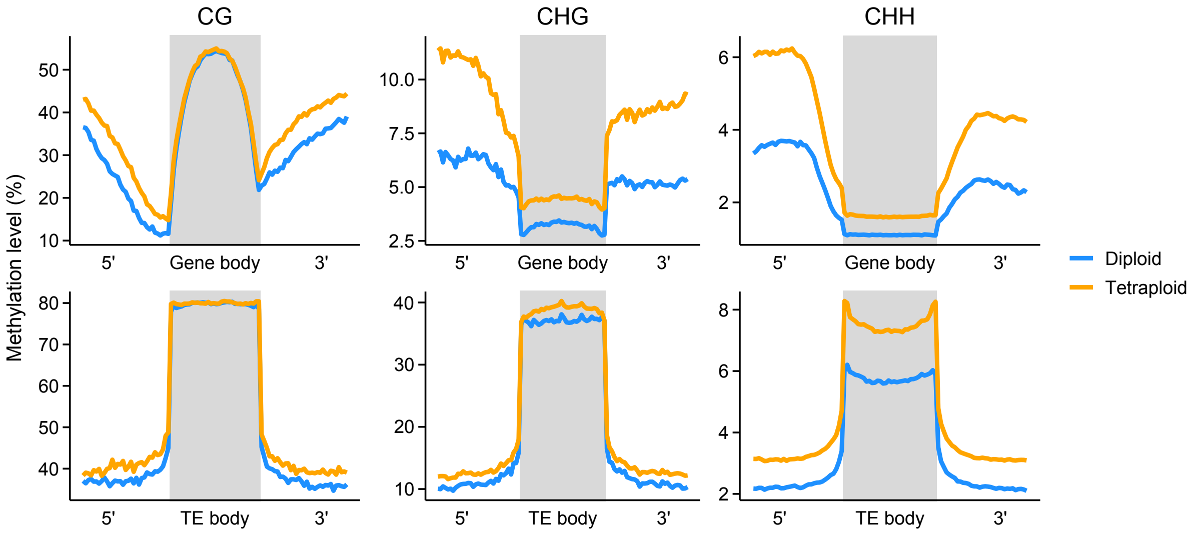


**Figure S4.** Methylation levels across meta-genes and meta-TEs, shown for CG, CHG, and CHH contexts. Note the difference in *y*-axis scales between the panels.

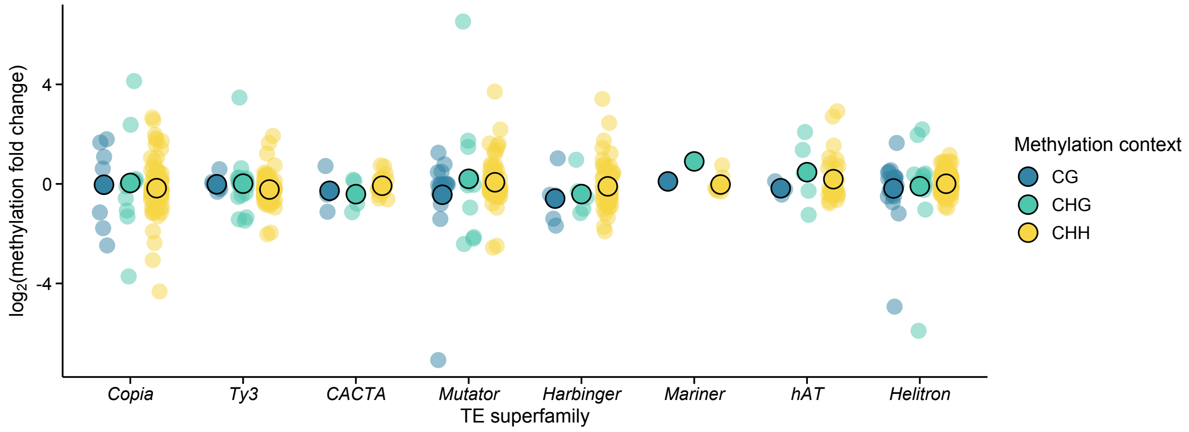


**Figure S5.** Methylation difference between diploids and tetraploids shown for differentially methylated TE families.

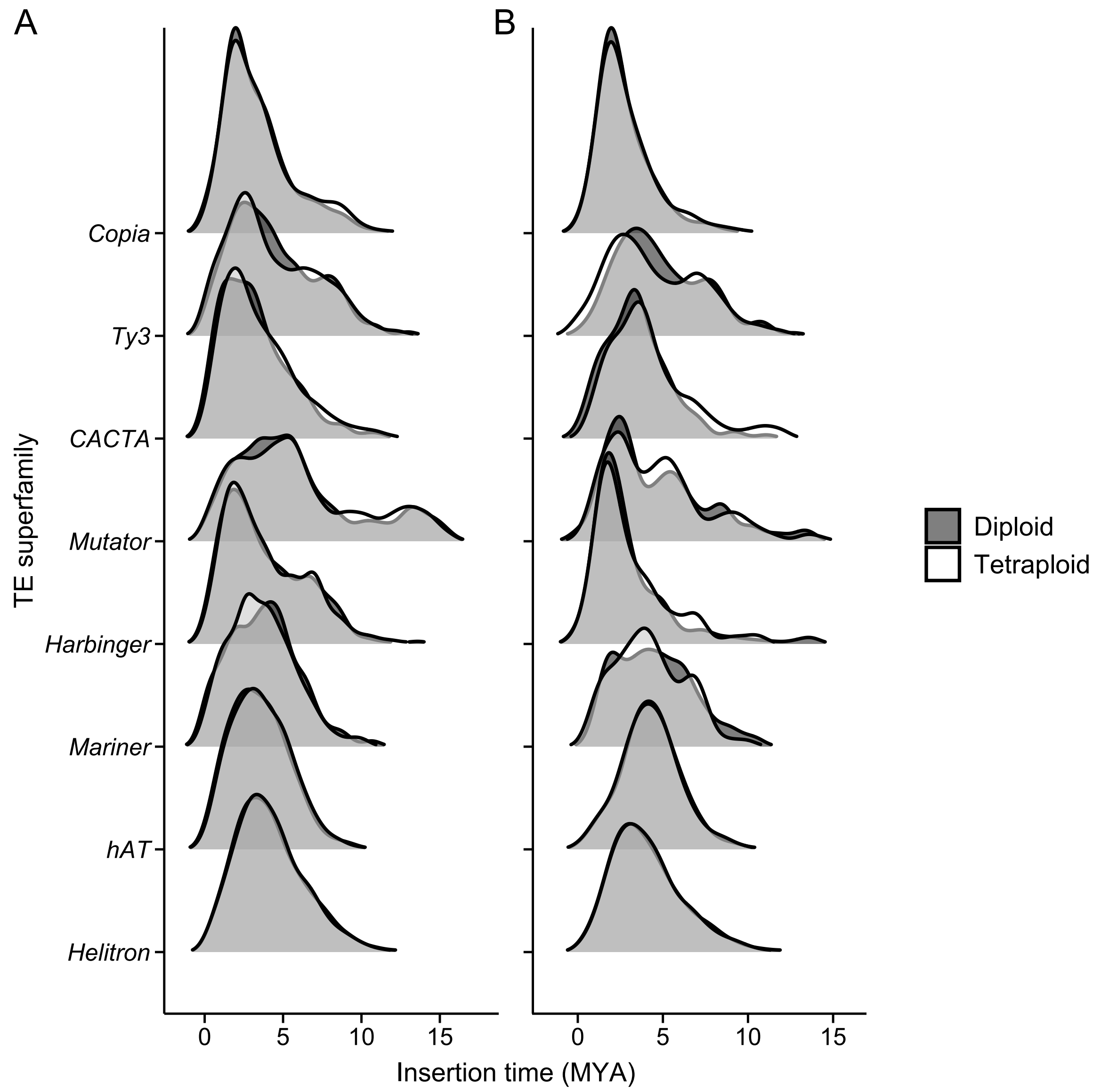


**Figure S6.** Distributions of TE insertions times in diploids and tetraploids. **A:** All TE families. **B**: Differentially methylated TE families.

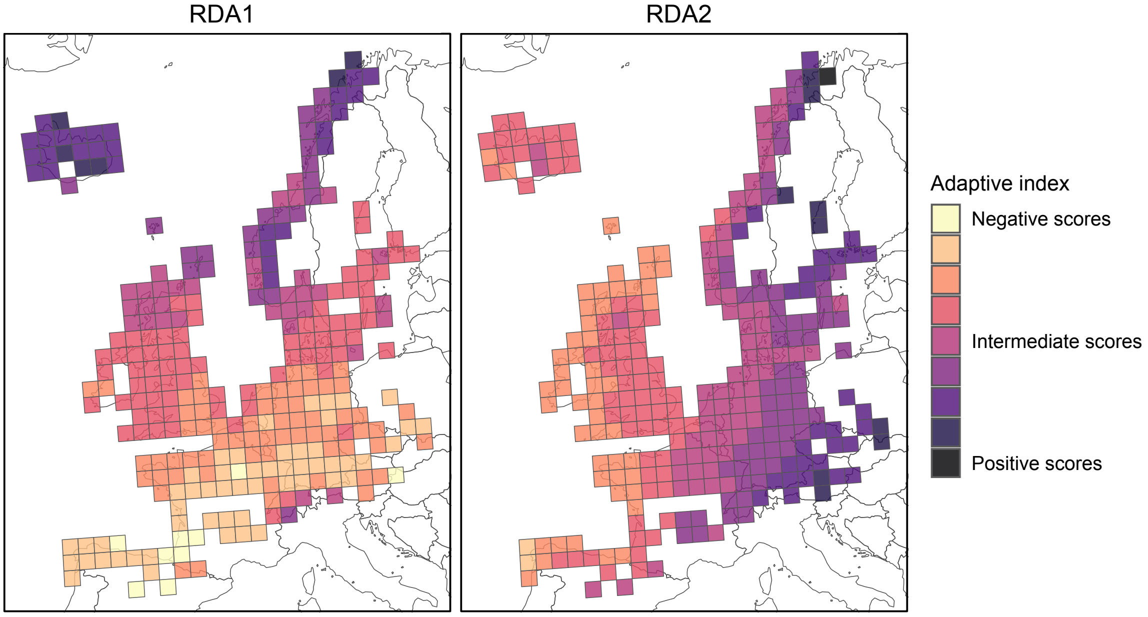


**Figure S7**. Adaptive index (based on combined SV and SNP outliers) predicted across the European range of *Cochlearia*. Similar colours indicate higher similarity in the genetic composition of the populations. Shown are two RDA axes used in the prediction of adaptive distance between SVs and SNPs.

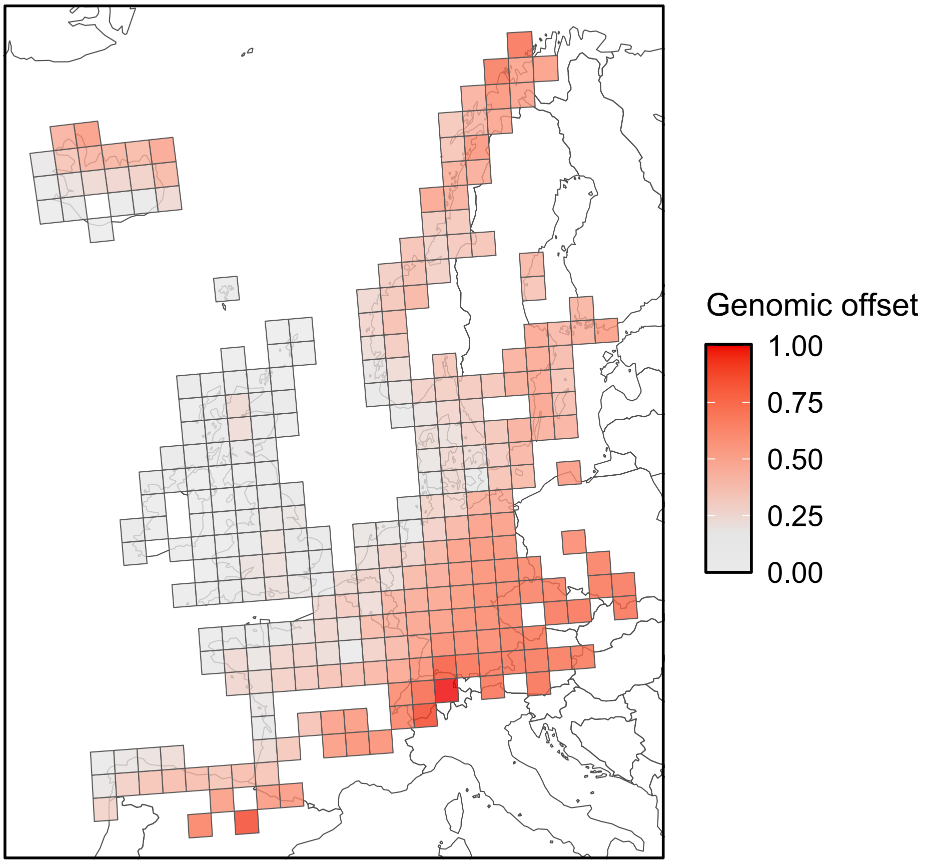


**Figure S8.** Genomic offset between current and future (2061 – 2080) climatic conditions, predicted using all climate-associated variants (SVs and SNPs). Colour scale indicates the relative level of genetic change that would be required to track climate change.

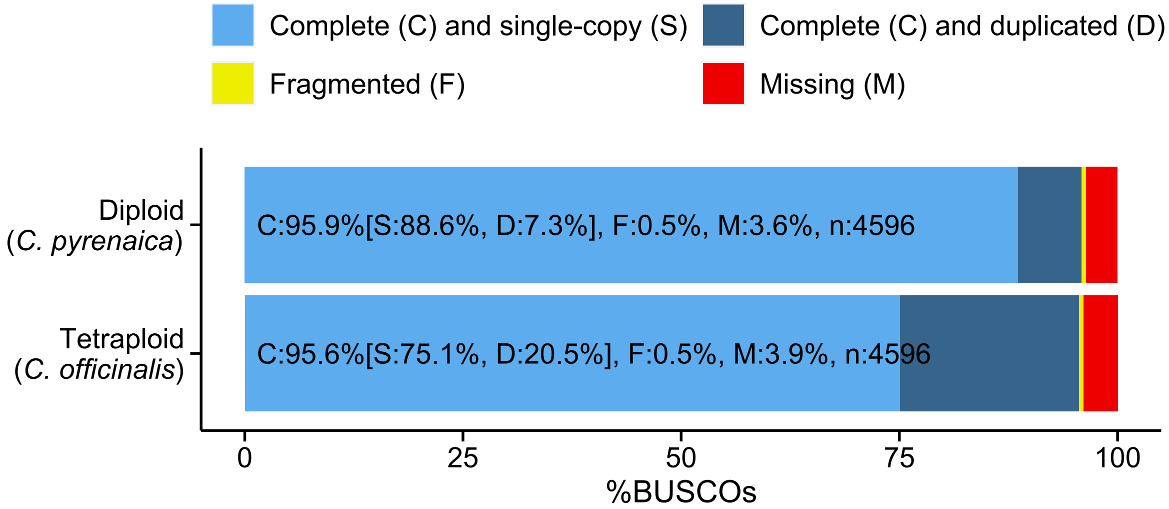


**Figure S9**. BUSCO results in the newly assembled diploid and tetraploid genomes.

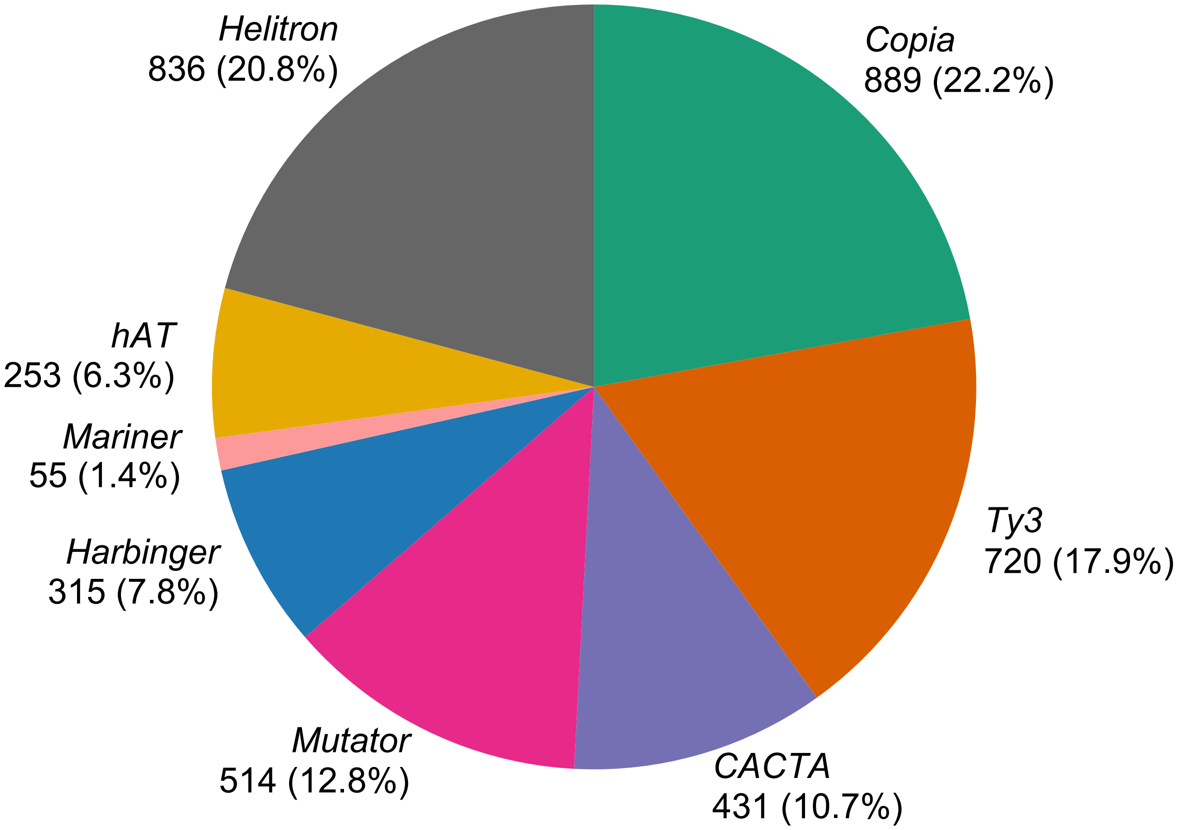


**Figure S10**. Number and proportion of TE superfamilies annotated in three different *Cochlearia* assemblies.

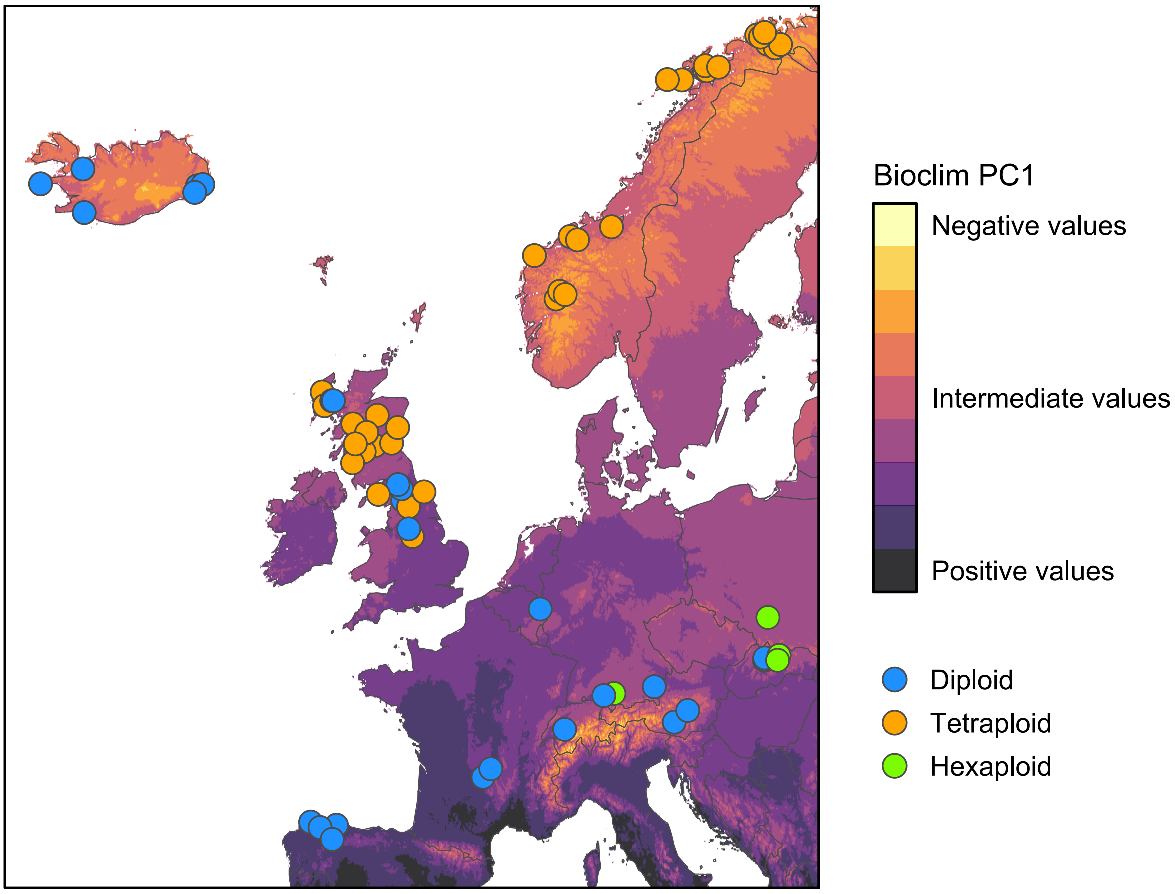


**Figure S11.** Geographical variation across 11 bioclimatic variables used in our GEA, summarised as the first principal component (PC) of the data. Coloured circles show 70 *Cochlearia* population used in the analyses. Following bioclim variables were used (in the order of their significance in explaining genetic variation among the populations):

bio2 = Mean diurnal range
bio3 = Isothermality
bio9 = Mean temperature of driest quarter
bio15 = Precipitation seasonality
bio18 = Precipitation of warmest quarter
bio8 = Mean temperature of wettest quarter
bio10 = Mean temperature of warmest quarter
bio7 = Temperature annual range
bio1 = Annual mean temperature
bio5 = Max temperature of warmest month
bio11 = Mean temperature of coldest quarter

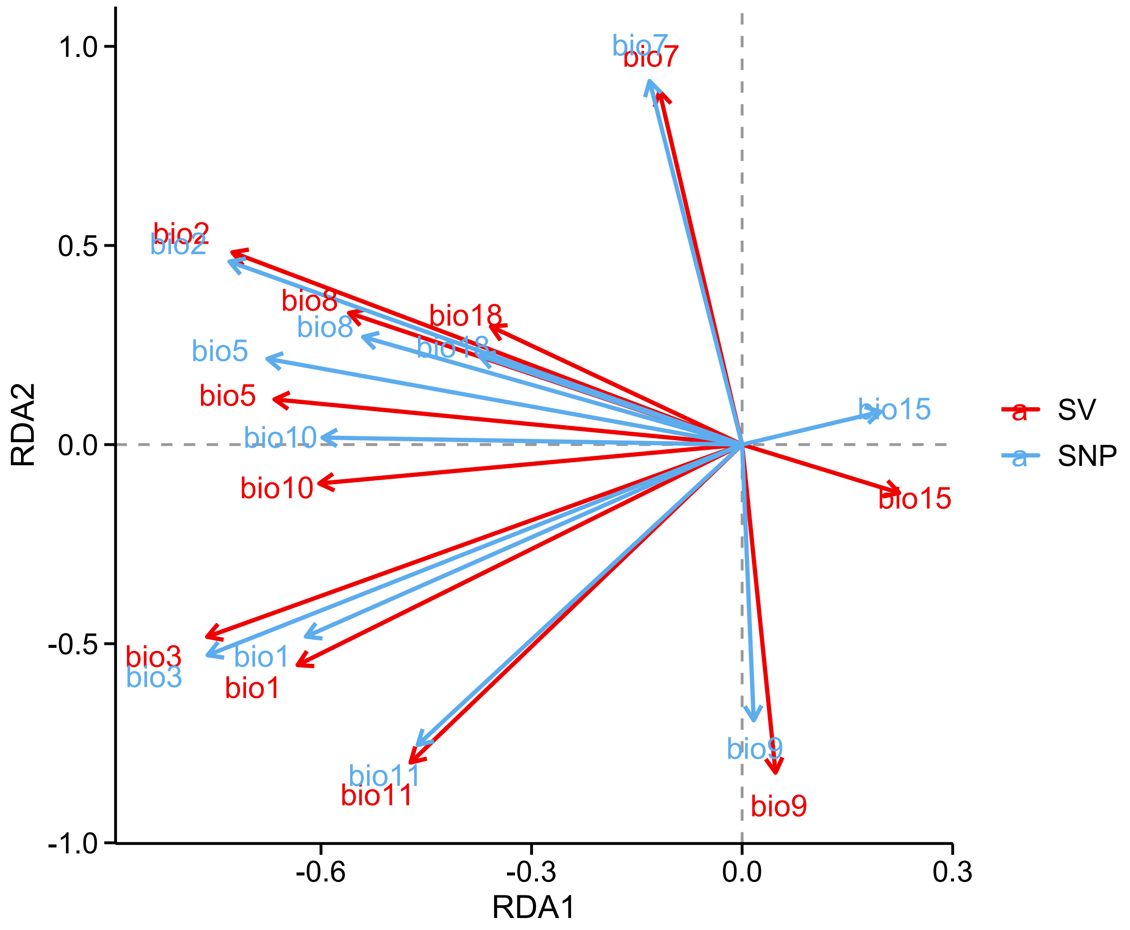


**Figure S12**. Loadings of 11 bioclimatic variables on first two RDA axes, shown for outlier SVs and SNPs. Length of the arrows indicate the load each of each variable on the axes.

**Table S1**. Summary of the long-read sequenced samples

| ID | Ploidy level | *Cochlearia* species | Country | Sequencing technology | Read length N50 (kb) | Mean depth |
|---|---|---|---|---|---|---|
| C10 | 2 | *pyrenaica* | OS | ONT | 8.9 | 26.1 |
| C12 | 2 | *pyrenaica* | DE | ONT | 5.8 | 24.1 |
| C13* | 2 | *pyrenaica* | DE | ONT | 8.7 | 9.0 |
| C14 | 2 | *aestuaria* | ES | ONT | 11.8 | 24.6 |
| C17 | 2 | *pyrenaica* | FR | ONT | 7.6 | 24.0 |
| C18* | 2 | *groenlandica* | IS | ONT | 4.3 | 9.1 |
| C19 | 2 | *aestuaria* | ES | ONT | 10.4 | 28.4 |
| C2 | 2 | *excelsa* | OS | ONT | 7.8 | 28.9 |
| C20 | 2 | *pyrenaica* | OS | ONT | 6.8 | 14.7 |
| C21 | 2 | *pyrenaica* | FR | ONT | 6.4 | 25.3 |
| C23* | 2 | *pyrenaica* | UK | ONT | 11.4 | 6.3 |
| C5* | 2 | *islandica* | IS | ONT | 10.3 | 8.2 |
| LAB 22 | 2 | *pyrenaica* | UK | ONT | 7.0 | 62.5 |
| NEN 30 | 2 | *pyrenaica* | UK | PacBio | 16.1 | 21.9 |
| BPS 1 | 4 | *officinalis* | UK | ONT | 7.3 | 82.6 |
| CUR | 4 | *officinalis* | UK | ONT | 5.4 | 48.7 |
| ELI 23 | 4 | *officinalis* | UK | ONT | 7.2 | 75.8 |
| ONI 12 | 4 | *officinalis* | UK | ONT | 7.6 | 58.1 |
| ROT 20 | 4 | *officinalis* | UK | ONT | 7.4 | 85.2 |
| ROT 26 | 4 | *officinalis* | UK | PacBio | 7.4 | 57.3 |
| TRE 13 | 4 | *officinalis* | UK | ONT | 8.5 | 69.4 |
| C27 | 6 | *tatrae* | SL | ONT | 5.8 | 18.3 |
| C24 | 8 | *anglica* | UK | ONT | 4.9 | 27.6 |

*Excluded from the main analyses due to low sequencing depth.

**Table S2**. Validation of Sniffles2 (v2.0.6) using simulated autotetraploids

| Sniffles mode | SV type | Mutation | Simulated depth | | | | |
|---|---|---|---|---|---|---|---|
| | | | 5× | 10× | 20× | 40× | 80× |
| Normal | Deletion | Simplex | 0.09 | 0.32 | 0.32 | 0.70 | 0.87 |
| Normal | Deletion | Duplex | 0.37 | 0.82 | 0.95 | 1.00 | 1.00 |
| Normal | Deletion | Triplex | 0.60 | 0.93 | 1.00 | 1.00 | 1.00 |
| Normal | Deletion | Quadruplex | 0.79 | 1.00 | 1.00 | 1.00 | 1.00 |
| Rare | Deletion | Simplex | 0.05 | 0.23 | 0.65 | 0.94 | 0.95 |
| Rare | Deletion | Duplex | 0.24 | 0.69 | 0.86 | 0.98 | 1.00 |
| Rare | Deletion | Triplex | 0.41 | 0.76 | 0.97 | 1.00 | 1.00 |
| Rare | Deletion | Quadruplex | 0.59 | 0.91 | 0.99 | 1.00 | 1.00 |
| Normal | Insertion | Simplex | 0.01 | 0.23 | 0.20 | 0.47 | 0.60 |
| Normal | Insertion | Duplex | 0.16 | 0.61 | 0.75 | 0.99 | 1.00 |
| Normal | Insertion | Triplex | 0.32 | 0.83 | 0.97 | 0.99 | 1.00 |
| Normal | Insertion | Quadruplex | 0.57 | 0.94 | 0.99 | 1.00 | 1.00 |
| Rare | Insertion | Simplex | 0.00 | 0.17 | 0.57 | 0.84 | 0.97 |
| Rare | Insertion | Duplex | 0.10 | 0.54 | 0.87 | 0.99 | 1.00 |
| Rare | Insertion | Triplex | 0.24 | 0.67 | 0.93 | 0.99 | 1.00 |
| Rare | Insertion | Quadruplex | 0.51 | 0.82 | 0.98 | 0.99 | 1.00 |

Shown are recall estimates for different levels of read depth over simulated insertions and deletions (100 repeats). For each of the simulated scenarios, precision estimates were > 0.95.

**Tables S3**. Species used in estimating coding sequence conservation with GERP++.

| Species | Source |
|---|---|
| *Aethionema arabicum* | https://plantcode.cup.uni-freiburg.de/aetar_db/ |
| *Arabidopsis thaliana* | Ensembl Plants |
| *Arabis alpina* | http://www.arabis-alpina.org/refseq.html |
| *Boechera stricta* | Phytozome |
| *Brassica rapa* | Ensembl Plants |
| *Camelina sativa* | Ensembl Plants |
| *Capsella rubella* | Phytozome |
| *Cardamine hirsuta* | http://chi.mpipz.mpg.de/assembly.html |
| *Carica papaya* | Phytozome |
| *Citrus clementina* | Ensembl Plants |
| *Cucumis sativus* | Ensembl Plants |
| *Draba nivalis* | https://doi.org/10.5061/dryad.pg4f4qrm4 |
| *Eucalyptus grandis* | Ensembl Plants |
| *Eutrema salsugineum* | Phytozome |
| *Fragaria vesca* | Phytozome |
| *Glycine max* | Ensembl Plants |
| *Gossypium raimondii* | Ensembl Plants |
| *Juglans regia* | Ensembl Plants |
| *Lobularia maritima* | https://ngdc.cncb.ac.cn/gwh/Assembly/9796/show |
| *Malus domestica* | Ensembl Plants |
| *Manihot esculenta* | Ensembl Plants |
| *Medicago truncatula* | Ensembl Plants |
| *Pistacia vera* | Ensembl Plants |
| *Populus trichocarpa* | Ensembl Plants |
| *Quercus lobata* | Ensembl Plants |
| *Raphanus sativus* | https://plantgarden.jp/ |
| *Schrenkiella parvula* | Phytozome |
| *Theobroma cacao* | Ensembl Plants |
| *Vitis vinifera* | Ensembl Plants |
